# babette: BEAUti 2, BEAST2 and Tracer for R

**DOI:** 10.1101/271866

**Authors:** Richèl J.C. Bilderbeek, Rampal S. Etienne

## Abstract

1. In the field of phylogenetics, BEAST2 is one of the most widely used software tools. It comes with the graphical user interfaces BEAUti 2, DensiTree and Tracer, to create BEAST2 configuration files and to interpret BEAST2’s output files. However, when many different alignments or model setups are required, a workflow of graphical user interfaces is cumbersome.
2. Here, we present a free, libre and open-source package, `babette`: ‘BEAUti 2, BEAST2 and Tracer for R’, for the R programming language. `babette` creates BEAST2 input files, runs BEAST2 and parses its results, all from an R function call.
3. We describe `babette`’s usage and the novel functionality it provides compared to the original tools and we give some examples.
4. As `babette` is designed to be of high quality and extendable, we conclude by describing the further development of the package.

**Samenvatting:**

1. In de fylogenetica is BEAST2 een van de meest gebruikte hulpprogramma’s. Het is gebundeld met de grafische gebruiksinterface BEAUti 2, DensiTree en Tracer, om BEAST2-configuratiebestanden te maken en om BEAST2-outputbestanden te interpreteren. Echter, als veel verschillende aligneringen of modelopzetten nodig zijn, is een werkvolgorde van meerdere grafische gebruiksinterfaces onhandig.
2. Hier presenteren we een gratis, vrij en open-source package, `babette`: ‘BEAUti 2, BEAST2 en Tracer voor R’, voor de programmeertaal R. `babette` schrijft BEAST2-configuratiebestanden, start BEAST2 and verwerkt de resultaten, alles met een enkele R functie-aanroep.
3. We beschrijven hoe `babette` te gebruiken is en de nieuwe mogelijkheden die het biedt vergeleken met de originele programma’s, aan de hand van enkele voorbeelden.
4. Omdat `babette` ontworpen is voor uitbreidbaarheid en hoge kwaliteit, sluiten we af met het beschrijven van de verdere ontwikkeling van dit package.

## 1 Introduction

Phylogenies are commonly used to explore evolutionary hypotheses. Not only can phylogenies show us how species (or other evolutionary units) are related to each other, but we can also estimate relevant parameters such as extinction and speciation rates from them. There are many phylogenetics tools available to obtain an estimate of the phylogeny of a given set of species. BEAST2 (Bouckaert *et al*. 2014) is one of the most widely used ones. It uses a Bayesian statistical framework to estimate the joint posterior distribution of phylogenies and model parameters, from one or more DNA, RNA or amino acid alignments (see figure 1 for an overview of the workflow).

**Figure 1.**
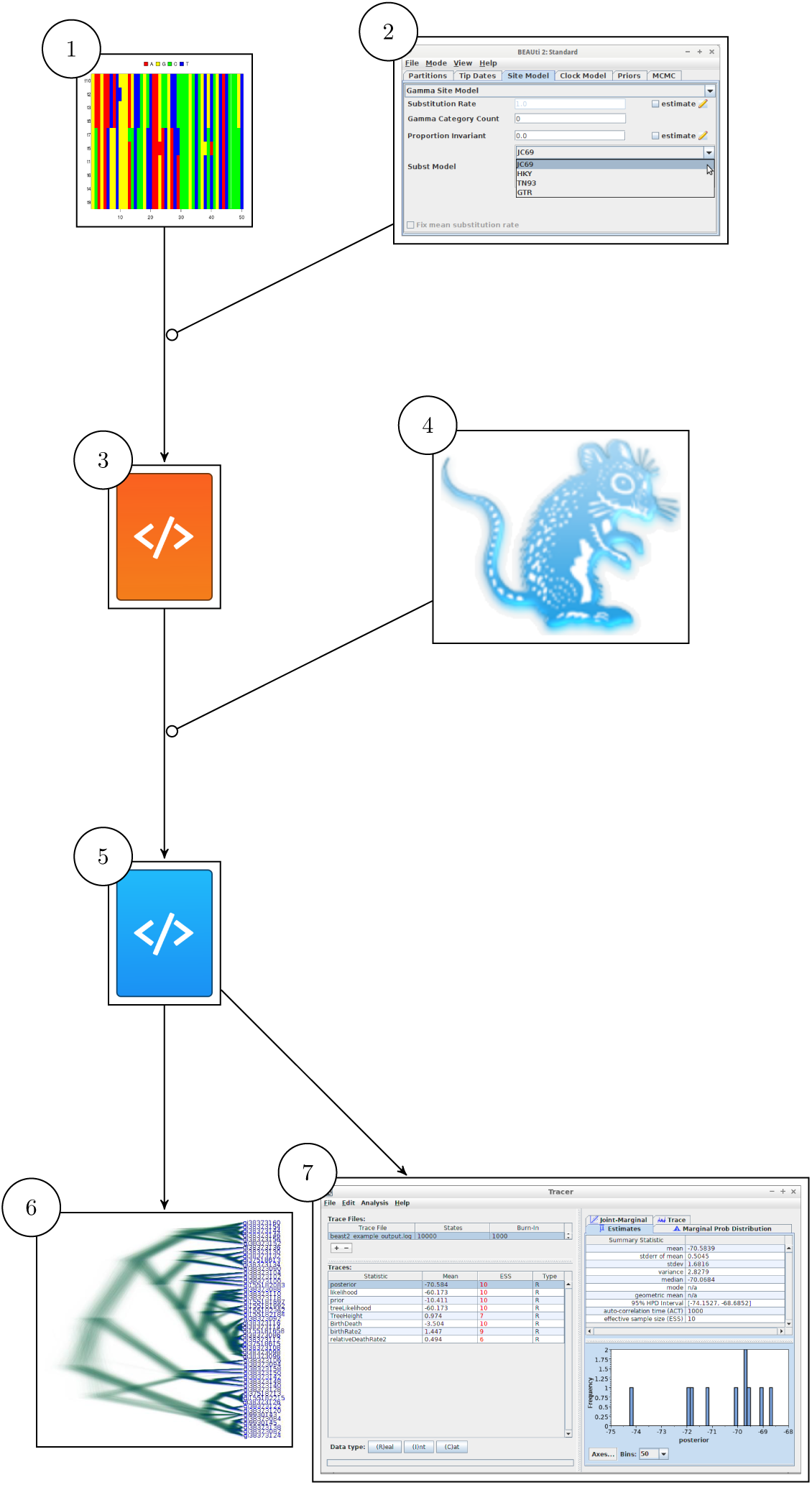

BEAST2 has a graphical and a command-line interface, that both need a configuration file containing alignments and model parameters. BEAST2 is bundled with BEAUti 2 (Drummond *et al*. 2012) (‘BEAUti’ from now on), a desktop application to create a BEAST2 configuration file. BEAUti has a user-friendly graphical user interface, with helpful default settings. As such, BEAUti is an attractive alternative to manual and error-prone editing of BEAST2 configuration files.

However, BEAUti cannot be called from a command-line script. This implies that when the user wants to explore the consequences of various settings, this must be done manually. This is the managable workflow when using a few alignments and doing a superficial analysis of sensitivity of the reconstructed tree to model settings. For exploring many trees (for instance from simulations), for a sliding-window analysis on a genomic alignment, or for a more thorough sensitivity analysis, one would like to loop through multiple (simulated or shortened) alignments, nucleotide substitution models, clock models and tree priors. One such tool to replace BEAUti is `BEASTmasteR` (Matzke 2015), which focuses on morphological traits and tip-dating, but also supports DNA data. `BEASTmasteR`, however, requires hundreds of lines of R code to setup the BEAST2 model configuration and a Microsoft Excel file to specify alignment files.

BEAST2 is also associated with Tracer (Rambaut & Drummond 2007) and DensiTree (Bouckaert & Heled 2014). Both are desktop applications to analyze the output of BEAST2, each with a user-friendly graphical user interface. Tracer’s purpose is to analyze the parameter estimates generated from a (BEAST1 and) BEAST2 run. It shows, among others, the effective sample size (ESS) and time series (‘the trace’, hence the name) of each variable in the MCMC run. Both ESS and trace are needed to assess the strength of the inference. DensiTree visualizes the phylogenies of a BEAST2 posterior, with many options to improve the simultaneous display of many phylogenies.

However, for exploring the output of many BEAST2 runs, one would like a script to collect all parameters’ ESSes, parameter traces and posterior phylogenies. There is no single package that offers a complete solution, but examples of R packages that offer a partial solution are rBEAST (Ratmann 2015) and RBeast (Faria & Suchard 2015). RBeast provides some plotting options and parsing of BEAST2 output files, but the plotting functions are too specific for general use. rBEAST was developed to test a particular biological hypothesis (Ratmann *et al*. 2016), and hence was not designed for general use.

Here, we present `babette`: BEAUti 2, BEAST2 and Tracer for R, which creates BEAST2 (v.2.4.7) configuration files, runs BEAST2, and analyzes its results, all from an R function call. This will save time, tedious mouse clicking and reduces the chances of errors in such repetitive actions. The interface of `babette` mimics the tools it is based on. This familiarity helps both beginner and experienced BEAST2 users to make the step from those tools to `babette`. `babette` enables the creation of a single-script pipeline from sequence alignments to posterior analysis in R.

## 2 Description

`babette` is written in the R programming language (R Core Team 2013) and enables the full BEAST2 workflow from a single R function call, in a similar way to what subsequent usage of BEAUti, DensiTree and Tracer would produce. `babette`’s main function is `bbt_run`, which configures BEAST2, runs it and parses its output. `bbt_run` needs at least the name of a FASTA file containing a DNA alignment. The default settings for the other arguments of `bbt_run` are identical to BEAUti’s and BEAST2’s default settings. Per alignment, a site model, clock model and tree prior can be chosen. Multiple alignments can be used, each with its own (unlinked) site model, clock model and tree prior.

`babette` currently has 108 exported functions to set up a BEAST2 configuration file. `babette` can currently handle the majority of BEAUti use cases. Because of BEAUti’s high number of plugins, `babette` uses a software architecture that is designed to be extended. Furthermore, `babette` has 13 exported functions to run and help run BEAST2. One function is used to run BEAST2, another one installs BEAST2 to a default location. Finally, `babette` has 21 exported function to parse the BEAST2 output files and analyze the created posterior. `babette` gives the same ESSes and summary statistics as Tracer. The data is formatted such that it can easily be visualized using `ggplot2` (for a trace, similar to Tracer) or `phangorn` (Schliep 2011) (for the phylogenies in a posterior, similar to DensiTree).

Currently, `babette` does not contain all functionality in BEAUti, BEAST2 and their many plug-ins, because these tools themselves also change in time. `babette` currently works only on DNA data, because this is the most common use case. Nevertheless, `babette` provides the majority of default tree priors and supports the most important command-line arguments of BEAST2, provides the core Tracer analysis options, and has the most basic subset of plotting options of DensiTree. Up till now, the `babette` features implemented are those requested by users. Further extension of `babette` will be based on future user requests.

## 3 Usage

`babette` can be installed easily from CRAN:

~~~
install . packages (“babette”)
~~~

For the most up-to-date version, one can download and install the package from `babette`’s GitHub repository:

~~~
devtools :: install_github (“richelbilderbeek/babette”)
~~~

To start using `babette`, load its functions in the global namespace first:

~~~
library (babette)
~~~

Because babette calls BEAST2, BEAST2 must be installed. This can be done from R, using:

~~~
install_beast2 ()
~~~

This will install BEAST2 to the default user data folder, but a different path can be specified as well. BEAUti, and likewise `babette`, needs at least a FASTA filename to produce a BEAST2 configuration file. In BEAUti, this is achieved by loading a FASTA file, then saving an output file using a common save file dialog. After this, BEAST2 needs to be applied to the created configuration file. It creates multiple files storing the posterior. These output files must be parsed by either Tracer or DensiTree. In `babette`, all this is achieved by:

~~~
out <- bbt_run (fasta_filenames = “anthus_aco.fas”)
~~~

This code will create a (temporary) BEAST2 configuration file, from the FASTA file with name `anthus_aco.fas` (which is supplied with the package, from Van Els & Norambuena 2018), using the same default settings as BEAUti, which are, among others, a Jukes-Cantor site model, a strict clock, and a Yule birth tree prior. `babette` will then execute BEAST2 using that file, and parses the output. The returned data structure, named `out`, is a list of parameter estimates (called `estimates`), posterior phylogenies (called `anthus_aco_trees`, named after the alignment’s name) and MCMC operator performance (`operators`). An example of using a different site model, clock model and tree prior is:

~~~
out <- bbt_run(
    fasta_filenames = “anthus_aco.fas”,
    site_models = create_hky_site_model (),
    clock_models = create_rln_clock_model (),
    tree_priors = create_bd_tree_prior ()
)
~~~

This code uses an HKY site model, a relaxed log-normal clock model and a birth-death tree prior, each with their default settings in BEAUti. Table 1 shows an overview of all functions to create site models, clock models and tree priors. Note that the arguments’ names `site_models`, `clock_models` and `tree_priors` are plural, as each of these can be (a list of) one or more elements. Each of these arguments must have the same number of elements, so that each alignment has its own site model, clock model and tree prior. An example of two alignments, each with its own site model, is:

~~~
out <- bbt_run (
    fasta_filenames = c (
        “anthus_aco . fas”,
        “anthus_nd2 . fas”
    ),
    site_models = list (
        create_tn93_site_model (),
        create_gtr_site_model ()
    )
)
~~~

**Table 1:**
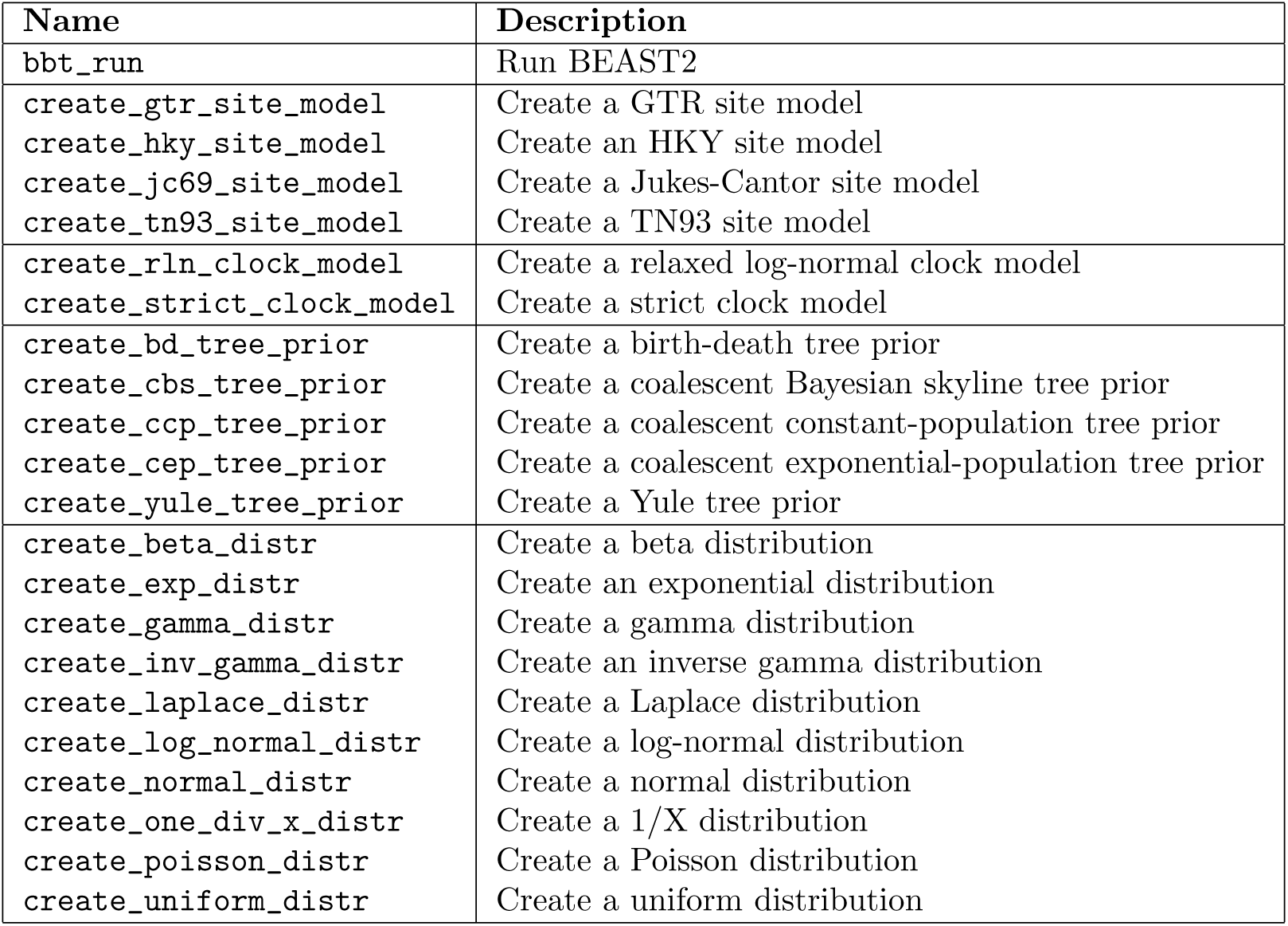
babette’s main functions

| Name | Description |
| --- | --- |
| bbt_run | Run BEAST2 |
| create_gtr_site_model | Create a GTR site model |
| create_hky_site_model | Create an HKY site model |
| create_jc69_site_model | Create a Jukes-Cantor site model |
| create_tn93_site_model | Create a TN93 site model |
| create_rln_clock_model | Create a relaxed log-normal clock model |
| create_strict_clock_model | Create a strict clock model |
| create_bd_tree_prior | Create a birth-death tree prior |
| create_cbs_tree_prior | Create a coalescent Bayesian skyline tree prior |
| create_ccp_tree_prior | Create a coalescent constant-population tree prior |
| create_cep_tree_prior | Create a coalescent exponential-population tree prior |
| create_yule_tree_prior | Create a Yule tree prior |
| create_beta_distr | Create a beta distribution |
| create_exp_distr | Create an exponential distribution |
| create_gamma_distr | Create a gamma distribution |
| create_inv_gamma_distr | Create an inverse gamma distribution |
| create_laplace_distr | Create a Laplace distribution |
| create_log_normal_distr | Create a log-normal distribution |
| create_normal_distr | Create a normal distribution |
| create_one_div_x_distr | Create a 1/X distribution |
| create_poisson_distr | Create a Poisson distribution |
| create_uniform_distr | Create a uniform distribution |

`babette` also uses the same default prior distributions as BEAUti for each of the site models, clock models and tree priors. For example, by default, a Yule tree prior assumes that the birth rate follows a uniform distribution, from minus infinity to plus infinity. One may prefer a different ddistribution instead. Here is an example how to specify an exponential distribution for the birth rate in a Yule tree prior in `babette`:

~~~
out <- bbt_run (
    fasta_filenames = “anthus_aco.fas”,
    tree_priors = create_yule_tree_prior (
        birth_rate_distr = create_exp_distr ()
    )
)
~~~

In this same example, one may specify the initial shape parameters of the exponential distribution. In BEAST2’s implementation, an exponential distribution has one shape parameter: its mean, which can be set to any value with `BEAUti`. To set the mean value of the exponential distribution to a fixed (non-estimated) value, do:

~~~
out <- bbt_run (
    fasta_filenames = “anthus_aco.fas”,
    tree_priors = create_yule_tree_prior (
        birth_rate_distr = create_exp_distr (
            mean = create_mean_param (
                value = 1.0,
                estimate = FALSE
            )
        )
    )
)
~~~

`babette` also supports node dating. Like BEAUti, one can specify Most Recent Common Ancestor (‘MRCA’) priors. An MRCA prior allows to specify taxa having a common ancestor, including a distribution for the date of that ancestor. With `babette`, this is achieved as follows:

~~~
out <- bbt_run (
    fasta_filenames = “anthus_aco.fas”,
    mrca_priors = create_mrca_prior (
      taxa_names = sample (get_taxa_names (“anthus_aco.fas”), size = 2),
      alignment_id = get_alignment_id (“anthus_aco.fas”),
      is_monophyletic = TRUE,
      mrca_distr = create_normal_distr (
        mean = create_mean_param (value = 15.0, estimate = FALSE),
        sigma = create_sigma_param (value = 0.025, estimate = FALSE)
      )
    )
)
~~~

Instead of dating the ancestor of two random taxa, any subset of taxa can be selected, and multiple sets are allowed. `babette` allows for the same core functionality as Tracer to show the values of the parameter estimates sampled in the BEAST2 run. This is called the “trace” (hence the name). The start of the trace, called the “burn-in”, is usually discarded, as an MCMC algorithm (such as used by BEAST2) first has to converge to its equilibrium and hence the parameter estimates are not representative. By default, Tracer discards the first 10% of all the parameter estimates. To remove a 20% burn-in from all parameter estimates in `babette`, the following code can be used:

~~~
traces <- remove_burn_ins (
    traces = out $ estimates,
    burn_in_fraction = 0.2
)
~~~

Tracer shows the ESSes of each posterior’s variables. These ESSes are important to determine the strength of the inference. As a rule of thumb, an ESS of 200 is acceptable for any parameter estimate. To calculate the effective sample sizes (of all estimated variables) in `babette`:

~~~
esses <- calc_esses (
    traces = traces,
    sample_interval = 1000
)
~~~

Tracer displays multiple summary statistics for each estimated variable: the mean and its standard error, standard deviation, variance, median, mode, geometric mean, 95% highest posterior density interval, auto-correlation time and effective sample size. It displays these statistics per variable. In `babette`, these summary statistics are collected for all estimated parameters at once:

~~~
sum_stats <- calc_summary_stats (
    traces = traces,
    sample_interval = 1000
)
~~~

`babette` allows for the same functionality as DensiTree. DensiTree displays the phylogenies in a posterior at the same time scale, drawn one over one another, allowing to see the uncertainty in topology and branch lengths. The posterior phylogenies are stored as `anthus_aco_trees` in the object `out`, and can be plotted as follows:

~~~
plot_densitree (phylos = out$anthus_aco_trees)
~~~

Instead of running the full pipeline, `babette` also allows to only create a BEAST2 configuration file. To create a BEAST2 configuration file, with all settings to default, use:

~~~
create_beast2_input_file (
    input_filenames = babette::get_babette_path (“anthus_aco.fas “),
    output_filename = “beast2.xml”
)
~~~

This file can then be loaded and edited by BEAUti, run by BEAST2, or run by `babette`:

~~~
run_beast2 (
    input_filename = “beast2.xml”,
    output_log_filename = “run.log”,
    output_trees_filenames = “posterior.trees”,
    output_state_filename = “final.xml.state”
)
~~~

`run_beast2` is a function that only runs BEAST2, and does not parse the output files (unlike `bbt_run`). In the example above, we specify the names of the desired BEAST2 output files explicitly, and these will be created in the R working directory, after which they can be inspected with other tools, or used to continue a BEAST2 run. When the names of these files are not specified, both `bbt_run` and `run_beast2` put these files in the default temporary folder (as obtained from `temp.dir()`) to keep the working directory clean of intermediate files.

## 4 babette resources

`babette` is free, libre and open source software available at http://github.com/richelbilderbeek/babette and is licensed under the GNU General Public License v3.0. `babette` uses the Travis CI (https://travis-ci.org) continuous integration service, which is known to significantly increase the number of bugs exposed (Vasilescu *et al*. 2015) and increases the speed at which new features are added (Vasilescu *et al*. 2015). `babette` has a 100% code coverage, which correlates with code quality (Horgan *et al*. 1994; Del Frate *et al*. 1995). `babette` follows Hadley Wickham’s style guide (Wickham 2015), which improves software quality (Fang 2001). `babette` depends on multiple packages, which are `ape` (Paradis *et al*. 2004), `beautier` (Bilderbeek 2018b), `beastier` (Bilderbeek 2018a), `devtools` (Wickham & Chang 2016), `geiger` (Harmon *et al*. 2008), `ggplot2` (Wickham 2009), `knitr` (Xie 2017), `phangorn` (Schliep 2011), `rmarkdown` (Allaire *et al*. 2017), `seqinr` (Charif & Lobry 2007), `stringr` (Wickham 2017), `testit` (Xie 2014) and `tracerer` (Bilderbeek 2018c). We tested `babette` to give a clean error message for incorrect input, by calling `babette` one million times with random or random sensible inputs, using a high performance computer cluster. The test scripts are supplied with `babette`.

`babette`’s development takes place on GitHub, https://github.com/richelbilderbeek/babette, which accommodates collaboration (Perez-Riverol *et al*. 2016) and improves transparency (Gorgolewski & Poldrack 2016). `babette`’s GitHub facilitates feature requests and has guidelines how to do so.

`babette`’s documentation is extensive. All functions are documented in the package’s internal documentation. For quick use, each exported function shows a minimal example. For easy exploration, each exported function’s documentation links to related functions. Additionally, `babette` has a vignette that demonstrates extensively how to use it. There is documentation on the GitHub to get started, with a dozen examples of BEAUti screenshots with equivalent `babette` code. Finally, `babette` has tutorial videos that can be downloaded or viewed on YouTube, https://goo.gl/weKaaU.

## 5 Citation of babette

Scientists using `babette` in a published paper can cite this article, and/or cite the `babette` package directly. To obtain this citation from within an R script, use:

~~~
> citation (“babette”)
~~~

## 6 Acknowledgements

Thanks to Yacine Ben Chehida and Paul van Els for supplying their BEAST2 use cases. Thanks again to Paul van Els for sharing his FASTA files for use by this package. Thanks to Leonel Herrera-Alsina, Raphael Scherrer and Giovanni Laudanno for their comments on this package and article. Thanks to Huw Ogilvie, Michael Matschiner and one anonymous reviewer for reviewing this article. Thanks to rOpenSci, and especially Noam Ross and Guangchuang Yu for reviewing the package’s source code. We would like to thank the Center for Information Technology of the University of Groningen for their support and for providing access to the Peregrine high performance computing cluster. We thank the Netherlands Organization for Scientific Research (NWO) for financial support through a VICI grant awarded to RSE.

## 7 Data Accessibility

All code is archived at http://github.com/richelbilderbeek/babette_article.

## 8 Authors’ contributions

RJCB and RSE conceived the idea for the package. RJCB created and tested the package, and wrote the first draft of the manuscript. RSE contributed substantially to revisions.

## References

Allaire, J., Xie, Y., McPherson, J., Luraschi, J., Ushey, K., Atkins, A., Wickham, H., Cheng, J. & Chang, W. (2017) rmarkdown: Dynamic Documents for R. R package version 1.8.

Bilderbeek, R.J. (2018a) beastier: BEAST2 from R. https://github.com/richelbilderbeek/beastier [Accessed: 2018–03–16].

Bilderbeek, R.J. (2018b) beautier: BEAUti 2 from R. https://github.com/richelbilderbeek/beautier [Accessed: 2018-03-16].

Bilderbeek, R.J. (2018c) tracerer: Tracer from R. https://github.com/richelbilderbeek/tracerer [Accessed: 2018-03-16].

Bouckaert, R. & Heled, J. (2014) Densitree 2: Seeing trees through the forest. bioRxiv, p. 012401.

Bouckaert, R., Heled, J., Kühnert, D., Vaughan, T., Wu, C.H., Xie, D., Suchard, M.A., Rambaut, A. & Drummond, A.J. (2014) Beast 2: a software platform for bayesian evolutionary analysis. PLoS Comput Biol, 10, e1003537.

Charif, D. & Lobry, J. (2007) SeqinR 1.0–2: a contributed package to the R project for statistical computing devoted to biological sequences retrieval and analysis. U. Bastolla, M. Porto, H. Roman & M. Vendruscolo, eds., Structural approaches to sequence evolution: Molecules, networks, populations, Biological and Medical Physics, Biomedical Engineering, pp. 207–232. Springer Verlag, New York. ISBN : 978–3–540–35305–8.

Del Frate, F., Garg, P., Mathur, A.P. & Pasquini, A. (1995) On the correlation between code coverage and software reliability. Software Reliability Engineering, 1995. Proceedings., Sixth International Symposium on, pp. 124–132. IEEE.

Drummond, A.J., Suchard, M.A., Xie, D. & Rambaut, A. (2012) Bayesian phylogenetics with beauti and the beast 1.7. Molecular biology and evolution, 29, 1969–1973.

Fang, X. (2001) Using a coding standard to improve program quality. Quality Software, 2001. Proceedings. Second Asia-Pacific Conference on, pp. 73–78. IEEE.

Faria, N. & Suchard, M.A. (2015) RBeast. https://github.com/beast-dev/RBeast [Accessed: 2018-03-02].

Gorgolewski, K.J. & Poldrack, R. (2016) A practical guide for improving transparency and reproducibility in neuroimaging research. bioRxiv, p. 039354.

Harmon, L., Weir, J., Brock, C., Glor, R. & Challenger, W. (2008) Geiger: investigating evolutionary radiations. Bioinformatics, 24, 129–131.

Horgan, J.R., London, S. & Lyu, M.R. (1994) Achieving software quality with testing coverage measures. Computer, 27, 60–69.

Matzke, N.J. (2015) BEASTmasteR: R tools for automated conversion of NEXUS data to BEAST2 XML format, for fossil tip-dating and other uses. https://github.com/nmatzke/BEASTmasteR [Accessed: 2018-02-28].

Paradis, E., Claude, J. & Strimmer, K. (2004) APE: analyses of phylogenetics and evolution in R language. Bioinformatics, 20, 289–290.

Perez-Riverol, Y., Gatto, L., Wang, R., Sachsenberg, T., Uszkoreit, J., Lepre-vost, F., Fufezan, C., Ternent, T., Eglen, S.J., Katz, D.S. et al. (2016) Ten simple rules for taking advantage of git and github. bioRxiv, p. 048744.

R Core Team (2013) R: A Language and Environment for Statistical Computing. R Foundation for Statistical Computing, Vienna, Austria.

Rambaut, A. & Drummond, A.J. (2007) Tracer v1.4. Available from http://beast.bio.ed.ac.uk/Tiacer.

Ratmann, O. (2015) rBEAST. https://github.com/olli0601/rBEAST [Accessed: 2018-03-02].

Ratmann, O., Van Sighem, A., Bezemer, D., Gavryushkina, A., Jurriaans, S., Wensing, A., De Wolf, F., Reiss, P., Fraser, C. et al. (2016) Sources of hiv infection among men having sex with men and implications for prevention. Science translational medicine, 8, 320ra2–320ra2.

Schliep, K. (2011) phangorn: phylogenetic analysis in R. Bioinformatics, 27, 592–593.

Van Els, P. & Norambuena, H.V. (2018) A revision of species limits in neotropical pipits anthus based on multilocus genetic and vocal data. Ibis.

Vasilescu, B., Yu, Y., Wang, H., Devanbu, P. & Filkov, V. (2015) Quality and productivity outcomes relating to continuous integration in github. Proceedings of the 2015 10th Joint Meeting on Foundations of Software Engineering, pp. 805–816. ACM.

Wickham, H. (2009) ggplot2: elegant graphics for data analysis. Springer New York.

Wickham, H. (2015) R packages: organize, test, document, and share your code. O’Reilly Media, Inc.

Wickham, H. (2017) stringr: Simple, Consistent Wrappers for Common String Operations. R package version 1.2.0.

Wickham, H. & Chang, W. (2016) devtools: Tools to Make Developing R Packages Easier. R package version 1.12.0.9000.

Xie, Y. (2014) testit: A Simple Package for Testing R Packages. R package version 0.4, http://CRAN.R-project.org/package=testit.

Xie, Y. (2017) knitr: A General-Purpose Package for Dynamic Report Generation in R. R package version 1.17.

